# CRISPR/Cas9 Treatment Causes Extended TP53-Dependent Cell Cycle Arrest In Human Cells

**DOI:** 10.1101/604538

**Authors:** Jonathan M. Geisinger, Tim Stearns

## Abstract

While the mechanism of CRISPR/Cas9 cleavage is understood, the large variation in mutant recovery for a given target sequence between cell lines is much less clear. We hypothesized that this variation may be due to differences in how the DNA damage response affects cell cycle progression. We used incorporation of EdU as a marker of cell cycle progression to analyze the response of several human cell lines to CRISPR/Cas9 treatment with a single guide directed to a unique locus. Cell lines with functionally wild-type TP53 exhibited higher levels of cell cycle arrest compared to lines without. Chemical inhibition of TP53 protein combined with *TP53* and *RB1* transcript silencing alleviated induced arrest in *TP53*^+/+^ cells. This arrest is driven in part by Cas9 binding to DNA. Additionally, wild-type Cas9 induced fewer 53BP1 foci in *TP53*^+/+^ cells compared to *TP53*^−/−^ cells, suggesting that differences in break sensing are responsible for cell cycle arrest variation. We conclude that CRISPR/Cas9 treatment induces a cell cycle arrest dependent on functional TP53 as well as Cas9 DNA binding and cleavage. Our findings suggest that transient inhibition of TP53 may increase genome editing efficiency in primary and *TP53*^+/+^ cell lines.

## INTRODUCTION

Genome engineering is a powerful tool, not only for modifying cells for therapeutic uses, but also for examining endogenous expression and localization of proteins, as well as aiding in deciphering their interactions with other proteins. In terms of semi-targeted insertion into the genome, the phage integrases, particularly phiC31, have been very useful for expressing exogenous cassettes in a variety of plants and animals for both translational and basic science purposes (reviewed in 1-2). However, targeting specific, single loci in the genome relies on homologous recombination (HR), which although highly efficient in yeast and some other simple model organisms (3–5), has been much less efficient in metazoans. With the development of the zinc-finger nucleases (ZFN; 6) and the TALE-nucleases (TALENs; 7), gene editing in metazoans and cells derived from them became a more feasible method of investigation and therapeutic development. These advances in the genome engineering toolbox built on seminal work with the I-SceI endonuclease (8), and greatly increased the efficiency of HR by introduction of a targeted double-strand break (DSB) in genomic DNA, while also allowing mutagenesis through the error-prone repair pathway due to the overhang-style of break induced by these nucleases.

Whereas the ZFNs and TALENs offered an unprecedented degree of control over where to induce a DSB due their modular, programmable nature, the design and cloning of these tools is challenging. This impediment has been reduced by discovery and application of the CRISPR/Cas9 system, which has proven to be a versatile tool for genome and transcriptome engineering. Consisting of a nuclease directed to a genomic target by a guide RNA, the engineered version of Cas9 requires only cloning the 20-base targeting sequence for the variant derived from *Streptococcus pyogenes*, for example (9–11). The major constraint on use of Cas9 is the requirement of a protospacer adjacent motif (PAM) downstream of the target sequence. This site varies in size and sequence between the different Cas9 proteins. For SpCas9, the PAM is canonically NGG (9), and the protein makes a predominantly blunt-ended DSB three to four bases 5’ from the PAM (12,13).

Whichever nuclease is used for genome engineering, there are still challenges faced in the efficient generation of the desired clones or organisms. For example, in addition to a rather cryptic target programming system, the ZFNs are highly cytotoxic (14). This cytotoxicity is greatly reduced when using TALENs, increasing the efficiency with which the desired modifications can be obtained (14). The cytotoxicity of the CRISPR/Cas9 varies between different cell lines, but is, unfortunately, high in induced pluripotent and embryonic stem cells (15,16), among the prime candidates for targeting. Also, the frequency of off-target mutations appears to be more severe with Cas9, as a recent report found larger deletions around the DSB and more complex rearrangements than previously identified (17). However, other studies, including both wild-type and nuclease-dead Cas9, have reported extremely high specificity, a virtual lack of true off-target activity, and an absence of complex rearrangements (18–20). Further compounding these issues is the high variance in gene editing efficiency between cell types and lines for a given guide RNA (13,21).

We asked if this high variance in gene editing efficiency could be explained by cell-line-specific cell cycle delay or arrest in response to CRISPR/Cas9 treatment. We reasoned that, because Cas9 predominantly generates blunt-ended DSBs (12,13), the DNA repair machinery would precisely repair the break through canonical NHEJ, which would enable Cas9 to target and cleave the same sequence again, enabling a cycle of damage and repair. This cycle would be expected to at least delay cells from progressing through the cell cycle and possibly arrest cells (22). Consequentially, this delay would reduce the efficiency of CRISPR/Cas9-based genome editing by reducing the frequency of modified cells, and the beneficial effect of removal of Cas9 from its target sequence via destabilization of Cas9 itself would be due to reducing this cell cycle delay. Such a destabilized Cas9 has been previously generated by fusing FKBP12-L106P destabilization domain (23) fused to the N-terminus of wild-type Cas9 to generate DD-Cas9 (13,24). DD-Cas9 can be inducibly and reversibly stabilized by the addition of the small molecule Shield-1 (13,23,24).

Here, we characterize the response of several human cell lines to wild-type, destabilization domain-linked, and nuclease-dead SpCas9 targeted to the safe harbour locus H11 on chromosome 22 (25). We identify a strong cell cycle delay/arrest phenotype that appears to be dependent on the TP53 status of each cell line. We determine that inhibition and transcript silencing of TP53 reduces the effect of this delay and illustrate that Cas9 appears to block the repair machinery from recognizing the DSB. Furthermore, we show that CRISPR/Cas9 treatment arrests cells to a higher degree than TALEN treatment.

## MATERIALS AND METHODS

### Cell Culture

hTERT RPE-1 wild-type and p53-null (from M-F. Tsou, Memorial Sloan Kettering Cancer Center) cells, HCT-116 wild-type and p53-null cells, HEK293T cells, and U2-OS cells were maintained in DMEM/F-12 media (Corning) supplemented with 10% Cosmic calf serum (HyClone Laboratories, Logan, UT). Cells were passaged with 0.25% Trypsin-EDTA (Corning) at a 1:10 split and maintained in 10 cm tissue culture dishes.

### Western Blotting

RPE1 cells were plated in 6-well plates one day prior were transfected with PEI and 2 μg of plasmid encoding the H11-r1-2 sgRNA and Cas9-DD or DD-Cas9 in the presence or absence of 0.5 μM Shield-1. After 24 hours, Shield-1 was washed out with PBS and replaced with fresh media. Protein was isolated at the time of washout, 24 hours later, and 48 hours later by extraction with RIPA buffer and immediately frozen.

For Western blotting using RPE1 cell lysate, 10 μg of protein were denatured in the presence of 1 x Laemmli buffer and 0.1 M DTT at 100 °C for 5 minutes. Samples were then resolved on an 8% SDS-PAGE gel at 100 V in 1x Running buffer. Protein was transferred to a nitrocellulose membrane (Bio-Rad) under wet transfer conditions at 200 mA for 1 hour with constant current at 4 °C. Following transfer, the membrane was blocked for 1 hour at room temperature with shaking in Tris-buffered saline + 0.2% TWEEN-20 (TBST) + 5% milk. Membranes were incubated overnight with agitation at 4 °C with 1:2000 monoclonal mouse anti-alpha-tubulin (Sigma-Aldrich; clone DM1A) and 1:1000 monoclonal mouse anti-Cas9 (Biolegend, San Diego, CA) The following day, membranes were washed 3 times for 5 minutes each with TBST. Secondary incubation was carried out at room temperature with agitation for 1 hour with either donkey anti-mouse IgG IRdye800CW (LI-COR Biosciences, Lincoln, NE) or goat anti-mouse IgG IRdye680RD (Li-Cor Biosciences) followed by washing 3 times for 5 minutes each with TBST. Membranes were then imaged on a LI-COR Odyssey (LI-COR Biosciences).

### Cell Cycle Progression Analysis

RPE1 cells were plated one day prior to transfection on poly-L-lysine-coated glass coverslips at 2.5 x 10^4^ cells per well in a 24-well plate. The following day cells were transfected via polyethylenimine with 500 ng of either pKER-Clover/pKER-NLS-Clover or a vector encoding the H11-r1-2 or H11 r2-3 sgRNA and WT-Cas9, DD-Cas9, Cas9-DD, or APEX2-X-dCas9, or a vector encoding wild-type SpCas9-P2A-mClover3, in the presence or absence of 0.5 μM Shield-1. 24 hours after transfection, Shield-1 was washed out with PBS and replaced with fresh media containing 10 μM EdU. Approximately 45 hours after Edu addition, cells were washed with cold PBS, fixed for 10 minutes at room temperature with 4% formaldehyde for 10 minutes, and rehydrated with PBS for 10 minutes. Cells were then blocked and permeabilized with PBS containing 5% bovine serum albumin and 0.2% Tween-20 (PBS-BT) for 30 minutes. After blocking, the click reaction between EdU and Alexafluor-594 azide (Life Technologies) was carried out using the Click-It 594 Labeling Kit (Life Technologies) according to the manufacturer’s instructions. After labeling and subsequent washing of cells three times for 5 minutes each with PBS + 0.2% Tween-20 (PBS-T), cells were incubated with 1:500 goat anti-GFP (Rockland Immunochemicals, Limerick, PA) and/or 1:1000 mouse IgG1 anti-Cas9 in PBSBT overnight at 4°C. The following day cells were washed three times with PBS-T for 5 minutes each and incubated for 1 hour in the dark at room temperature with DAPI and either 1:1000 donkey anti-goat IgG Alexafluor-488 (Life Technologies) or 1:1000 Donkey anti-mouse IgG1 Alexafluor-488 or 1:1000 Donkey anti-goat IgG Alexafluor-568 (Life Technologies). Coverslips were then washed with PBS three times for 5 minutes each and affixed to slides via MOWIOL.

Slides were photographed on an Axioskop 200M microscope. Three 423 μm x 324 μm windows were analyzed per coverslip in ImageJ by counting total number of nuclei, number of GFP+/Cas9+ cells, and number of GFP+/Cas9+ EdU+ nuclei and the average was calculated for each coverslip.

For flow cytometry-based analysis, cells were plated at 5×10^4^ to 7.5×10^4^ cells per well in 24-well plates and transfected 24 hours later as described above. Where appropriate, cells were treated with 20 μM pifithrin-α from 24 hours pre-transfection through the end of the experiment, with replacement every 24 hours. Shield-1, where appropriate, and EdU were added as described above. Roughly 45 hours post-EdU-addition, cells were harvested via trypsinization into 1.75 mL microcentrifuge tubes, centrifuged at 300 x g for 5 minutes at 4°C, washed with 800 μL PBS, centrifuged at 300 x g for 5 minutes at 4°C, and fixed for 10 minutes with 200 μL 4% formaldehyde. All subsequent centrifugations were carried out at 300 x g for 5 minutes at 4°C. Cells were then washed twice with 800 μL PBS before blocking and permeabilization as described above. Click chemistry and immunofluorescence were carried out as a described above with the following changes: anti-Cas9 was used at 1:500, all stainings were carried out for 30 minutes at room temperature in the dark, two washes were carried out between click chemistry, primary staining, and secondary staining with addition of 500 μL of PBS-T followed by centrifugation at 300 x g for 5 minutes at 4°C. Following the final wash after secondary staining, cell pellets were resuspended in 200 μL of PBS containing diluted DAPI (Sigma-Aldrich) and transferred to polypropylene FACS tubes for analysis.

For experiments involving short hairpin RNAs, the list of transfected vectors also included vectors expressing nuclear-localized Clover, WT-Cas9, and either a single short-hairpin expression cassette directed against *TP53* or two short-hairpin expression cassettes directed against *TP53* and *RB1*. The *RB1* short hairpin was cloned from pMKO.1 puro RB shRNA, a gift from William Hahn (Addgene plasmid # 10670). The *TP53* hairpin was cloned from pCXLE-hOCT3/4-shp53-F, a gift from Shinya Yamanaka (Addgene plasmid # 27077).

### Flow Cytometry

Flow cytometric analysis for this project was done on instruments in the Stanford Shared FACS Facility. Flow cytometric analysis was carried out on an LSRII-class analyzer (BD Biosciences, San Jose, CA).

### Double-Strand Break Visualization

3.5×10^4^, 5×10^4^, or 7.5×10^4^ HEK293T, WT-, or *TP53*^−/−^ RPE-1 cells were plated on poly-D-lysine-coated coverslips in a 24-well plate. 24 hours later, cells were transfected with vectors encoding nuclear-localized Clover and either WT-Cas9 and H11-r2-3 guide or DD-Cas9 and H11-r1-2 guide in the presence of 0.5 μM Shield-1 (which was washed out 24 hours post-transfection). At 3 days post-transfection, cells were washed with PBS, fixed with methanol for 10 minutes at −20 °C, rehydrated in cold PBS at room temperature for 10 minutes, and blocked with PBS-BT for a minimum of 30 minutes. Cells were then incubated with the following primary antibodies for 1 hour at room temperature: 1:500 mouse IgG2b anti-53BP1 (BD Biosciences) and 1:200 rabbit anti-p21 (eBioscience). Cells were then washed three times for 5 minutes each with PBS-T before secondary antibody staining in PBS-BT with the following antibodies and DAPI: 1:1000 Alexa488 donkey anti-goat IgG, 1:1000 Alexa568 donkey anti-mouse IgG, 1:1000 Alexa680 donkey anti-rabbit IgG. Cells were then washed three times for 5 minutes each with PBS-T, sealed to a slide with MOWIOL, and cured overnight. Slides were imaged on a Keyence BZX-710 fluorescence microscope (Keyence, Osaka, Japan). Cells were then quantified by counting transfected cells categorized by the presence or absence of 53BP1 foci and the presence of nuclear p21 in ImageJ.

### AraC-Based Proliferation Assay

RPE-1 WT and p53-null cells were plated at 1.25 x10^5^ cells per well in each well of a 12-well tissue culture plate. The following day, cells were transfected in triplicate with PEI and vectors encoding nuclear-localized Clover alone or WT-Cas9 and H11r2-3 guide, one of a pair of TALENs targeting the human H11 locus, or both TALENs. 24 hours after transfection, ara-C (Sigma-Aldrich) was added to 2 wells of each condition at a concentration of 100 μM. The third replicate of each condition was treated with a volume of water equivalent to the volume of ara-C added to the media. Ara-C treatment was maintained for 5 days with replacement every 2 days. After 5 days of treatment, the cells were imaged on an IncuCyte Zoom (Essen BioScience, Ann Arbor, Michigan), with 9 images taken per well. The IncuCyte’s analysis software was then used to calculate the number of transfected cells per image. Afterwards, the ratio of surviving Clover+ cells in each ara-C replicate relative to the water control for that condition was calculated and compared to a theoretical ratio of 1 using a one-sample t-test.

### Statistics

All statistical analysis was carried out using Graphpad Prism 7 for Windows (Graphpad Software, Inc.).

## RESULTS

### Wild-Type SpCas9 Leads to Dilution of Transfected Cells in the Population by Limiting S-Phase Progression in RPE-1 Cells

Having previously characterized the behaviour of destabilized variants of SpCas9 in HEK293T cells (13), we sought to characterize the kinetics of destabilized SpCas9 in a more primary-like immortalized cell line, RPE-1, with the goal of optimizing genome engineering in these cells. In this experiment, we transfected RPE-1 cells with the DD-Cas9 and WT-Cas9 vectors in the presence or absence of the stabilizing small molecule ligand Shield-1 for 24 hours before washout. We examined three time points: immediately after washout (24 hours post-transfection), 4 hours post-washout, and 24 hours post-washout. Western blotting revealed little change in DD-Cas9 levels by 24 hours post-washout, regardless of Shield-1 treatment, as would be expected due to continued production of the protein and its large size (Supplementary Figure S1A). However, we observed that WT-Cas9 level decreased more rapidly than the destabilization variants by 24 hours post-washout, which is consistent with dilution of transfected cells within the population (Supplementary Figure S1A).

We remained concerned about the apparent dilution of cells transfected with wild-type Cas9 as compared to the DD-Cas9-transfected cells, thus choosing to examine localization of these proteins. We reasoned that this dilution could be caused by delayed cell cycle progression or cell cycle arrest generated from repeated cycles of cleavage and repair by Cas9. To examine the localization of stabilized and destabilized Cas9, we transfected RPE-1 cells with a vector encoding the H11-r1-2 sgRNA and WT-Cas9, or DD-Cas9, in the presence or absence of Shield-1 for 24 hours with and without washout. pKER-Clover (13), a vector encoding Clover fluorescent protein under the control of the *EF1α* promoter, was used as a transfection control. To examine the effect of CRISPR/Cas9 on cell cycle progression, we labeled newly-synthesized DNA in transfected cells with EdU for approximately two doublings (45-48 hours) following Shield-1 washout. We then visualized EdU-containing DNA and Cas9. Fluorescence microscopy revealed that WT-Cas9 and stabilized DD-Cas9 localized throughout the nucleus and cytoplasm, whereas DD-Cas9 in the absence of stabilization appeared reduced in protein level and post-washout DD-Cas9 appeared mainly as cytoplasmic puncta (Supplementary Figure S1B). The percent transfected cells was similar between treatments (Supplementary Figure S1C). However, there was a striking difference between treatments in the percent of cycling cells within the Cas9+ population as visualized by EdU incorporation (Supplementary Figure S1D). While stabilized DD-Cas9 post-washout (43.11 ± 4.80% cycling cells in the Cas9+ population) displayed a significant difference compared to stabilized DD-Cas9 (*P* = 0.0022), the most significant difference was observed in comparison to WT-Cas9 (7.77 ± 2.33%; *P* = 0.0001), indicating a severe delay of cell cycle progression associated with WT-Cas9. This CRISPR/Cas9-associated cell cycle delay was even more striking when visualized as percentage of total nuclei, which indicated that EdU+ Cas9+ nuclei made up only 0.94 ± 0.37 % of the total number of nuclei (Supplementary Figure S1E). Interestingly, we did not detect a significant difference in cycling cells between stabilized DD-Cas9 post-washout and pKER-Clover, suggesting that washout of Shield-1 truly leads to destabilization of DD-Cas9 (Supplementary Figure S1D; *P* = 0.0803). These results indicated removal of Shield-1 led to cytoplasmic accumulation of DD-Cas9 and alleviated CRISPR/Cas9-associated cell cycle arrest. Also, these results suggest that the decrease in WT-Cas9 protein levels observed by Western blot in RPE-1 cells is due to dilution of the signal by actively dividing non-transfected cells.

### CRISPR/Cas9-Associated Cell Cycle Delay is Highly Variable Between Different Cell Types

Given the low level of EdU+ cells observed in the CRISPR/Cas9-treated cells and the relatively low-throughput of our immunofluorescence approach, we repeated our experiment in wild-type RPE-1 cells using flow cytometry. Additionally, we included HEK293T, U2-OS, *TP53*^−/−^ RPE-1, and wild-type and *TP53*^−/−^ HCT-116 cells in our panel. To control for transfection efficiency, we included a nuclear-localized Clover (NLS-Clover) reporter cassette on the CRISPR/Cas9 vector under the control of the *EF1α* promoter (Figure 1A). We followed the same experimental timeline as we used for the immunofluorescence analysis (Figure 1B).

**Figure 1.**
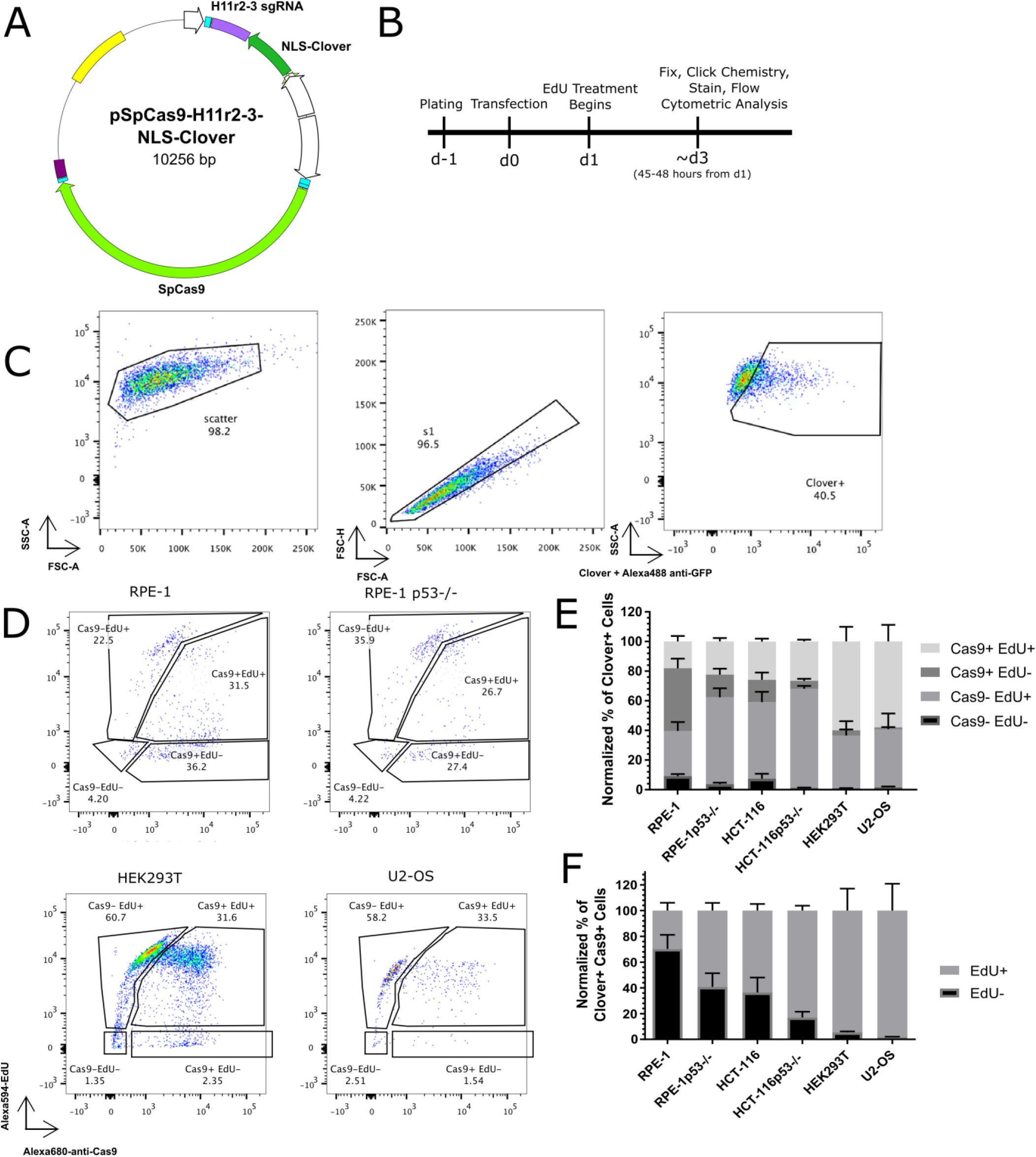
CRISPR/Cas9 treatment decreases cell cycle progression. (A) Schematic of transfected plasmid, identifying the H11 r2-3 gRNA cassette, the NLS-Clover cassette, and the WT-Cas9 cassette. (B) Experimental design schematic. (C) Flow cytometry gating strategy for identifying the transfected (Clover+) subpopulation. (D) Representative flow cytometric data of transfected cells for Cas9 and EdU positivity for the indicated cell lines. (E) Bar graph of the normalized percentages of four Cas9 EdU subpopulations in transfected cells for the six indicated cell lines. *n* = at least two independent experiments consisting of three technical replicates for each transfection. Data are shown as the mean ± SEM. (F) Bar graph of the Cas9+ normalized percentages of EdU+ and EdU-from panel E. Data is shown as the mean ± SEM.

After gating on Clover+ cells (Figure 1C), we further analyzed cells based on Cas9 and EdU positivity (Figure 1D). We first categorized the Clover+ population into [Cas9+ EdU+], [Cas9+ EdU-], [Cas9-EdU+], and [Cas9-EdU-] subpopulations, and noted that transfected HEK293T and U2-OS cells displayed striking larger percentages of Cas9+ EdU+ cells than both types of RPE-1 and HCT-116 cells (Figure 1E). We then focused our analysis on the Cas9+ subpopulations, reasoning that transfected cells expressing Cas9 at detectable levels would be most likely to be cells possessing DSBs (Figure 1F). After normalization of the Cas9+ population, we observed that WT RPE-1 cells displayed the lowest percentage of EdU+ cells (29.78 ± 6.12 % of Cas9+ cells), whereas HEK293T and U2-OS cells displayed the highest amounts of EdU+ cells (94.51 ± 17.01 and 98.18 ± 20.88 %, respectively). We also observed that the *TP53*^−/−^ RPE-1 and HCT-116 cell lines displayed higher levels of EdU+ cells than their wild-type counterparts. These results indicated that the flow cytometry-based assay is more amenable to high-throughput analysis, that cell cycle progression is inhibited to varying degrees in response to CRISPR/Cas9 treatment in different cell types, and that TP53 mediates this inhibition of cell cycle progression.

### Mediators of CRISPR/Cas9-Associated Cell Cycle Arrest

Our comparison of cell cycle progression in different cell lines suggested that *TP53* status may mediate cell cycle arrest in response to CRISPR/Cas9 treatment, similar to the conclusion reached by other groups (16,26). We then investigated the effect of abrogating TP53 functionality in wild-type RPE-1 cells. We took a two-prolonged approach of transcript knockdown via the addition of a short-hairpin cassette against *TP53* to our WT-Cas9 vector targeting the H11 locus (Figure 2A) and functional protein inhibition via the small molecule pifithrin-α (27, Figure 2B). We considered additional factors affecting cell cycle progression in CRISPR/Cas9-treated cells based on the cell lines used. Because U2-OS cells are reported to lack functional RB1 (28), which is activated in response to DNA damage (29), we constructed a WT-Cas9-expression vector containing a short-hairpin expression cassette targeting *RB1* in addition to the *TP53*-targeting hairpin. We theorized that that the combination of transcript knockdown and functional protein inhibition would alleviate CRISPR/Cas9-inudced cell cycle arrest. To test this idea, we used our flow cytometry-based assay, with the only changes being the pretreatment of cells with pifithrin-α 24 hours before transfection and the presence of pifithrin-α throughout the experiment (Figure 2C). We found that the only the combination of pifithrin-α and both short hairpin cassettes significantly increased the EdU+ subpopulation of the transfected Cas9+ cells (an increase of 18.54 ± 5.87%; *P* = 0.011). Additionally, we noted that treatment of wild-type RPE-1 cells with pifithrin-α and the TP53 short hairpin cassette resulted in similar levels of CRISPR/Cas9-associated cell cycle arrest as untreated p53-null RPE-1 cells (Supplementary Figure S2). However, combined treatment or treatment with only the *TP53* short hairpin increased CRISPR/Cas9-associated cell cycle arrest in p53-null RPE-1 cells (Supplementary Figure S2; *P* = 0.0282 and *P* = 0.254, respectively), which suggests the presence of a low-affinity, off-target transcript only revealed in the absence of *TP53* message. These results demonstrate that CRISPR/Cas9-associated cell cycle arrest can be partially alleviated by inhibiting TP53.

**Figure 2.**
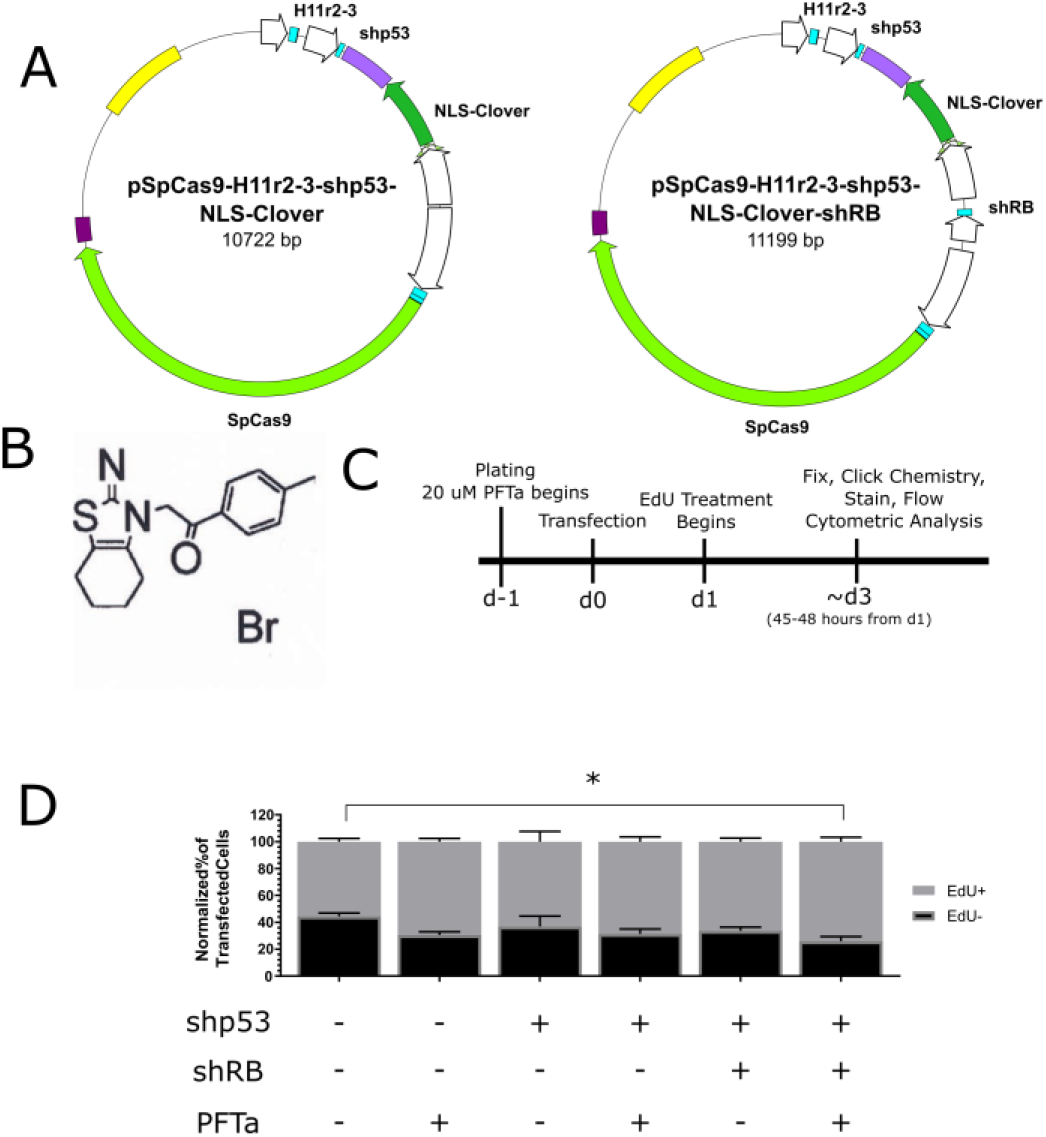
Inhibition of TP53 can alleviate CRISPR/Cas9-associated cell cycle arrest. (A) Schematic of Cas9 reporter plasmids containing a short-hairpin cassette against *TP53* or two short hairpin cassettes against *TP53* and *RB1*. (B) Chemical structure of pifithrin-α. (C) Schematic of experimental design. (D) Bar graph of Cas9+ normalized percentages of EdU+ and EdU-cells for indicated treatments. *n* = at least two independent experiments each consisting of three technical replicates for each transfection. Data is shown as the mean ± SEM. * = *P* < 0.05. Data were analyzed via an ordinary two-way ANOVA followed by a post-hoc Dunnett’s multiple comparisons test, where each treatment was compared to wild-type RPE-1 cells treated with wild-type Cas9.

### CRISPR/Cas9-Associated Cell Cycle Arrest is Dependent on Targeting, Cleavage, and the Presence of a Guide RNA

Having investigated the possibility of alleviating CRISPR/Cas9-associated cell cycle arrest, we next sought to identify the molecular mechanism of the arrest. To this end, we reasoned that there are two non-exclusive mechanisms of action: (1) cleavage by Cas9, and (2) targeting and binding DNA by Cas9. We investigated these possibilities using the EdU-based flow cytometry assay with transfected cells.

To address if cleavage by Cas9 contributed to CRISPR/Cas9-associated cell cycle arrest, we chose to investigate the cell cycle response to targeted nuclease-dead Cas9 (dCas9) using our EdU-based flow cytometry assay. To this end, we repurposed an APEX2-tagged variant of dCas9 (AX-dCas9) that we had originally constructed for labelling and proteome mapping of genomic loci, much like the C-BERST approach (30). We then transfected HEK293T, U2-OS, wild-type and *TP53*^−/−^ HCT-116, and wild-type and *TP53*^−/−^ RPE-1 cells with a vector encoding NLS-Clover fluorescent protein, AX-dCas9, and a guide RNA cassette targeting the H11 locus (Figure 3A). We observed no difference in population proportions for HEK293T and U2-OS cells, as expected from our experiments with WT-Cas9. However, we observed an increase in the percent of transfected cycling cells lacking detectable AX-dCas9 in *TP53*^−/−^ RPE-1 cells (75.33 ± 4.25%) compared to wild-type RPE-1 cells (32.75 ± 8.48%), indicative of Cas9 binding to DNA contributing to cell cycle arrest in a TP53-dependent manner and the existence of a threshold for Cas9 detection in our assay (Figure 3B). We also noted that AX-dCas9-transfected *TP53*^−/−^ RPE-1 cells displayed lower total levels of Cas9+ cells compared to their WT-Cas9-transfected counterparts, illustrative of a stronger effect of cleavage and binding on cell cycle arrest compared to binding alone. In examining the Cas9+ fraction of transfected cells, we again observed that *TP53*-deficient cell lines displayed higher levels of EdU+ cells compared to their wild-type counterparts (Figure 3C). Taken together, these results reinforce the role of TP53 in mediating CRISPR/Cas9-associated cell cycle arrest and identify that the binding of Cas9 to DNA can lead to cell cycle arrest in a TP53-dependent manner.

**Figure 3.**
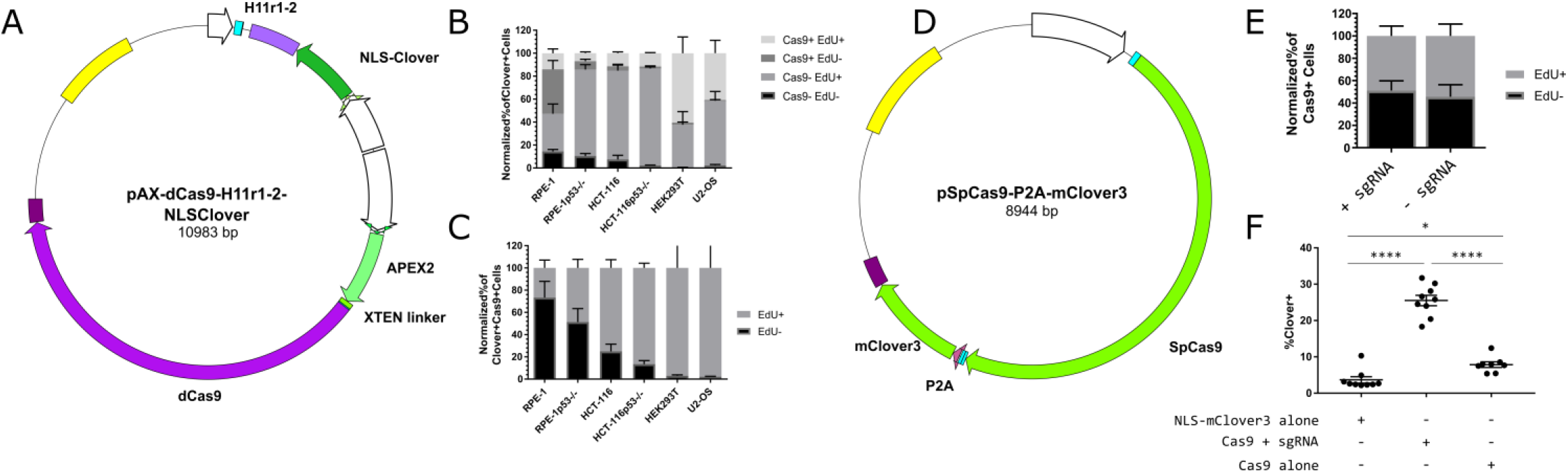
CRISPR/Cas9-associated cell cycle arrest is mediated by Cas9 binding. (A) Schematic of APEX2-dCas9 plasmid encoding the H11 r1-2 gRNA and an NLS-Clover cassette. (B) Bar graph of the Clover+-normalized percentages of four Cas9 EdU subpopulations in AX-dCas9-transfected cells for the six indicated cell lines. *n* = at least two independent experiments consisting of three technical replicates for each transfection. Data are shown as the mean ± SEM. (C) Bar graph of the Cas9+ normalized percentages of EdU+ and EdU-from panel B. Data is shown as the mean ± SEM. (D) Schematic of plasmid encoding WT-Cas9-P2A-mClover3 without a gRNA cassette. (E) Bar graph of Cas9+ normalized percentages of EdU+ and EdU-transfected wild-type RPE-1 cells with and without the H11r2-3 gRNA. *n* = three independent experiments consisting of at least two technical replicates each. Data are shown as the mean ± SEM. (F) Dot plot of percentage of Clover+ cells for indicated transfections. *n* = three independent experiments consisting of at least two technical replicates each. Bars indicate mean ± SEM. * = *P* < 0.05, **** = *P* < 0.0001. Data were analyzed via a one-way ANOVA followed by a post-hoc Sidak’s multiple comparisons test.

To test whether cell cycle arrest is dependent on targeting and binding of Cas9 to genomic DNA, we generated a WT-Cas9-expressing vector lacking a guide RNA expression cassette, but possessing mClover3, a brighter, optimized variant of Clover (31), under the control of the CAG promoter and separated from Cas9 by a P2A skipping peptide sequence (Figure 3D). We compared wild-type RPE-1 cells transfected with this vector or the H11-targeting WT-Cas9 vector and found no difference in the Cas9+ population for EdU incorporation (Figure 3E). However, upon examining the percentage of cells that were fluorescent-protein+ three days after transfection in our flow cytometry-based assay, we observed that the fluorescence protein+ percentage of Cas9-alone treated cells was more similar to Clover-alone transfected cells than to Cas9+sgRNA-treated cells (Figure 3F; a difference of 4.20 ± 1.53% [*P* = 0.0316] versus 17.65 ± 1.53% [*P* < 0.0001]). This result indicates not only that targeted binding of Cas9 to genomic DNA contributes to CRISPR/Cas9-associated cell cycle arrest, but that Cas9 protein or the Cas9 expression vector may also contribute to cell cycle arrest. However, it is unclear if the latter specifically contributes to CRISPR/Cas9-associated cell cycle arrest.

### Bound CRISPR/Cas9 Complex Blocks DSB Recognition

Having identified that CRISPR/Cas9-associated cell cycle arrest is dependent on both binding to and cleavage of DNA by Cas9, we hypothesized that Cas9 may prevent DNA damage recognition proteins, such as 53BP1, from detecting resulting DSB. Such a hypothesis is attractive because of Cas9’s exceptionally long DNA occupancy time (32) and recent evidence that high levels of transcription through the targeted area may lead to higher levels of editing, presumably through dislodging of Cas9 from the target DNA (33). Additionally, we reasoned that the presence of 53BP1 foci would also indicate a degree of cell cycle progression (34).

To these ends, we adopted a similar experimental design as for the flow-cytometry-based assays, but we used immunofluorescence-based microscopy to assay cells at the end point of 3 days post-transfection (Figure 4A). Additionally, we used DD-Cas9 (13) as a way to theoretically remove Cas9 from the DNA and expose the DSB. We modified the DD-Cas9 vector by inserting a self-contained nuclear-localized Clover expression cassette.

**Figure 4.**
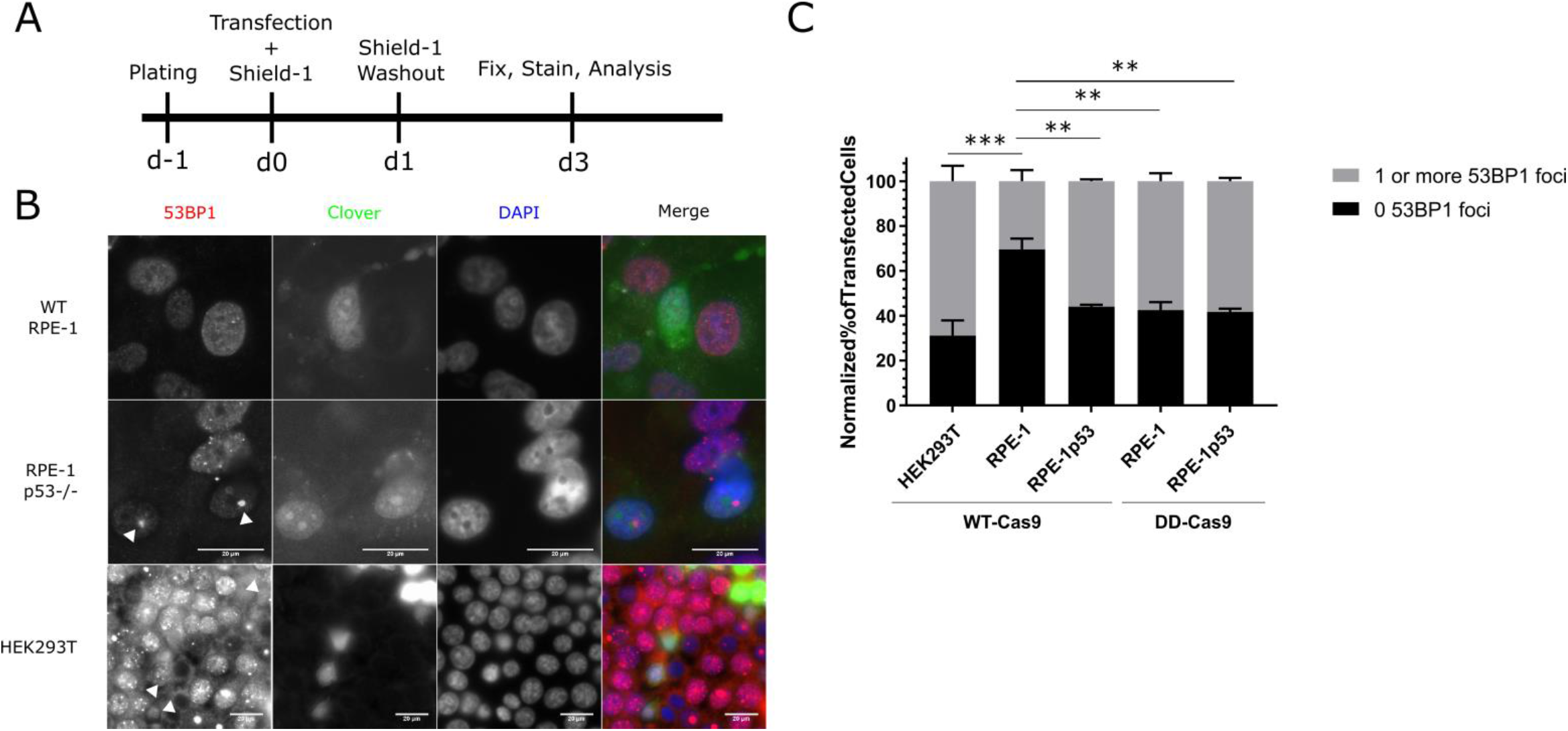
Wild-type Cas9 blocks recognition of Cas9-induced double-strand breaks. (A) Experimental design schematic. (B) Representative immunofluorescence of transfected wild-type and *TP53*^−/−^ RPE-1 and HEK293T cells for 53BP1 foci. White arrowheads indicate Clover+ cells possessing one or more 53BP1 foci. Scale bar indicates 20 μm. (C) Bar graph illustrating percentages of given cell lines transfected with the indicated vector that possessed either zero 53BP1 foci or one or more 53BP1 foci. *n* = at least two independent experiments consisting of at least 60 transfected cells per experiment. Data are shown as the mean ± SEM. *** = *P* < 0.001, **** = P = 0.0001. Data were analyzed via an ordinary two-way ANOVA followed by a post-hoc Dunnett’s multiple comparisons test, where each treatment was compared to wild-type RPE-1 cells treated with wild-type Cas9.

To examine whether the cell is capable of recognizing and repairing Cas9-induced DSBs, we transfected HEK293T, wild-type RPE-1, and TP53^−/−^ RPE-1 cells with vectors encoding a nuclear-localized Clover expression cassette, a guide RNA expression cassette targeting the H11 locus, and either WT-Cas9 or DD-Cas9 in the presence of Shield-1 for 24 hours before washout before examining the cells via immunofluorescent microscopy at roughly 3 days post-transfection (Figure 4A). We then identified transfected cells by Clover/GFP immunofluorescence and categorized cells by the presence or absence of 53BP1 foci within the nucleus (Figure 4B, data shown for wild-type Cas9). Additionally, we examined cells for significant levels of nuclear p21, which would indicate canonical TP53-mediated cell cycle arrest. We observed that wild-type RPE-1 cells transfected with wild-type Cas9 displayed the highest level of cells lacking 53BP1 foci, whereas *TP53*^−/−^ RPE-1 cells transfected with wild-type Cas9 displayed higher levels of cells possessing one or more 53BP1 foci (Figure 4C). We additionally observed that the choice of DD-Cas9 had no difference in *TP53*^−/−^ RPE-1 cells, but resulted in a significantly higher percentage of cells possessing one or more 53BP1 foci in wild-type RPE-1 cells compared to wild-type Cas9 treatment (57.54 ± 3.57% versus 3± 4.93%; *P* = 0.0033). Consistent with the results from the cell cycle progression assay, HEK293T cells transfected with wild-type Cas9 displayed the highest percentage of cells with 53BP1 foci (68.92 ± 6.81%). Also, we observed virtually no transfected cells with levels of nuclear p21 that would be consistent with canonical TP53-mediated arrest. These results demonstrate that wild-type Cas9 prevents recognition of nuclease-induced DSBs, and that this effect can be ameliorated either by elimination of TP53, or by destabilization of Cas9.

Because of the effect of destabilizing DD-Cas9 on DSB recognition, we sought to characterize the effect of DD-Cas9 on cell cycle progression using the EdU-based flow cytometry assay. We transfected HEK293T, U2-OS, wild-type and *TP53*^−/−^ HCT-116, and wild-type and *TP53*^−/−^ RPE-1 cells with the DD-Cas9 vector described above. In *TP53*^−/−^ RPE-1 cells, we observed an increase in the level of transfected, cycling cells lacking detectable Cas9 compared to WT RPE-1 cells, consistent with the effect of TP53 on CRISPR/Cas9-associated cell cycle arrest and, at least for *TP53*^−/−^ RPE-1, consistent with destabilization of DD-Cas9 after washout (Supplementary Figure S3A). However, the population proportions of DD-Cas9-treated WT RPE-1 cells appeared unchanged from those of WT-Cas9-treated RPE-1 cells (Figure 1E), consistent with cleavage leading to cell cycle delay or arrest. In examining the Cas9+ fraction of transfected cells, we once more observed that *TP53*-deficient cell lines displayed higher levels of EdU+ cells compared to their wild-type counterparts (Supplementary Figure S3B). Taken together, these results further support the role of TP53 in mediating CRISPR/Cas9-associated cell cycle arrest and are consistent with the recent report that DSBs generated by DD-Cas9 lead to cell cycle delay even after repair (35).

### CRISPR/Cas9 Treatment Arrests Cells to a Greater Extent Than TALEN Treatment

Having identified a cell cycle progression defect in CRISPR/Cas9-treated cells through lack of EdU incorporation, we chose to further investigate if this was truly an arrest phenotype. At the same time, we wanted to determine if this cell cycle progression defect is unique to the CRISPR/Cas9 system. To this end, we modified vectors encoding a pair of TALENs targeted against the human H11 locus (25) with a self-contained nuclear-localized mClover3 expression cassette (Figure 5A).

**Figure 5.**
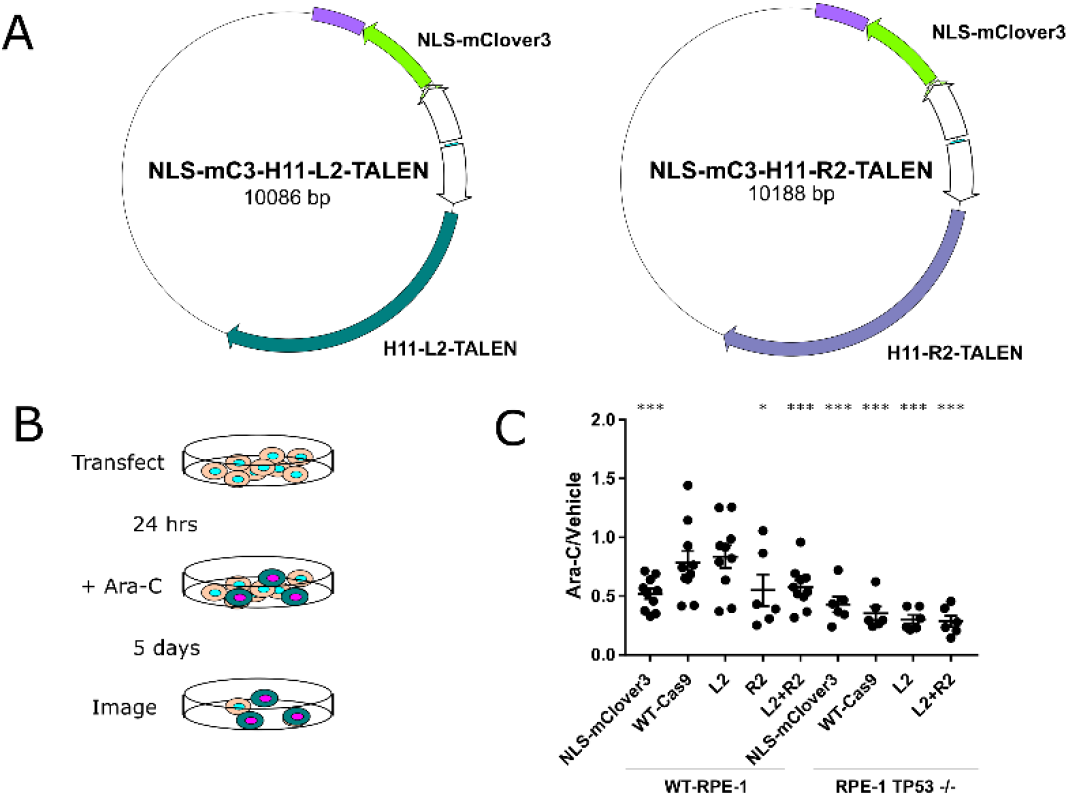
CRISPR/Cas9 treatment leads to a greater degree of cell cycle arrest than TALEN treatment. (A) Schematic of H11 L2 and R2 TALEN plasmids encoding NLS-mClover3. (B) Schematic of experimental design of ara-C-based cell cycle arrest assay. (C) Dot plot of ratios of Clover+ cells after 5 days of ara-C treatment versus vehicle for indicated cell lines and transfected vectors. *n* = three independent experiments each consisting of two ara-C-treated technical replicates and one vehicle-treated control. Error bars indicate SEM. * = *P* < 0.05, *** = *P* < 0.0005. Data were analyzed with a one-sample t-test comparing the actual mean against a theoretical mean of 1.

For the experimental design, we transfected wild-type and *TP53*^−/−^ RPE-1 cells with (1) a vector encoding a nuclear-localized mClover3, (2) a vector encoding WT-Cas9 targeted against the H11 locus and a nuclear-localized Clover expression cassette, (3 and 4) either of the previously described TALEN plasmids, or (5) both H11 TALEN plasmids 24 hours after plating. After 24 hours, we began treating the cells with 20 μM cytosine arabinoside (ara-C), a cytosine analog that kills cells actively synthesizing DNA, for 5 days, at which point we quantified surviving transfected cells using an Incucyte Zoom live cell analyser (Figure 5B). We then used the Incucyte’s software to automate counting of Clover+ cells in each image to compare control-treated to ara-C-treated wells. In our analysis, we calculated the ratio of Clover+ cells in ara-C treatment to control, reasoning that a ratio not significantly different from 1 would indicate a degree of cell cycle arrest in the transfected cells, whereas ratios significantly different than 1 would indicate cell cycle progression in the case where the ratio is less than 1. We observed that *TP53*^−/−^ RPE-1 cells, regardless of vector, displayed significant cell cycle progression, whereas the wild-type RPE-1 cells displayed significant cell cycle progression only in the presence of mClover3 alone, both TALENs together, and the R2 TALEN alone, but not for targeted wild-type Cas9 or the L2 TALEN alone (Figure 5C). These results indicate that TALEN treatment leads to less cell cycle arrest than CRISPR/Cas9 treatment, and underscores the requirement of *TP53* in mediating this arrest. Additionally, these results combined with the extended duration of ara-C treatment provide further evidence that CRISPR/Cas9-treated cells do indeed undergo a cell cycle arrest.

## DISCUSSION

In this work, we find that CRISPR/Cas9 treatment leads to extended cell cycle arrest in some human cell lines. This arrest requires, in part, functional TP53 and RB1, and can be alleviated to a degree by manipulation of TP53 activity. Additionally, we found that this arrest is mediated by DNA cleavage, DNA binding, and the Cas9 protein itself, in descending order of importance. Furthermore, we provide evidence that the propensity of Cas9 to remain bound to DNA after cleavage reduces the ability of cells to recognize and repair the double-strand break, and that this too is dependent on functional TP53.

Our work demonstrating an extended cell cycle arrest in response to CRISPR/Cas9 treatment has several implications for the planning, conduct, and interpretation of gene editing and genome engineering experiments. First, and consistent with other recent studies (16,26,35,36,37), our work suggests that caution is required in the interpretation of CRISPR/Cas9-based screens. Currently, the outcomes of such screens are usually assessed by the retention/loss of guides targeting specific genes from larger pools of guides, and are interpreted as identifying the sufficiency or necessity of these specific genes for the phenotype in question (38). Our work raises the possibility of such screens unintentionally selecting for pre-existing cells with non-functional TP53 or for cells possessing pre-existing mutations in the target locus, rendering the target locus uneditable. Such a possibility would bias results of the screen, leading to false positives or false negatives, depending on the nature of the screen. Selection of pre-existing mutants might be acceptable in fulfilling a screen, unless one is interested specifically in alleles derived from Cas9-mediated cutting. Second, our work identifies TP53 inhibition as a potential means to increase desired clone recovery in directed genome engineering CRISPR/Cas9 experiments. This inhibition can accomplished in several ways, including mutation of *TP53*, treatment with a small-molecule inhibitor, and/or transcript silencing by RNAi. Given that even temporary inhibition of TP53 can result in an increase genome instability, an alternative is to use an inducible destabilized Cas9, which somewhat mitigates the detrimental effect of Cas9 expression (13,24,35). Finally, our work raises the question of what, exactly, does Cas9 treatment is selecting for in the variety of experiments in which it used. Our results suggest that investigators should take into account that CRISPR/Cas9 treatment is likely selecting for cells that are untargetable due to pre-existing endogenous variation at that locus, and thus are somewhat resistant to the growth inhibition effect of Cas9 treatment.

Our results are largely consistent with two recently-published works that also identified TP53 as being important in both a CRISPR/Cas9-induced DNA damage response and an induced cytotoxicity response (16,26). In the work of Haapaniemi and colleagues (26), TP53 was identified as mediating a DNA-damage response in RPE-1 cells; the screen they performed also identified GATA6, CDKN1A/P21, and RB1. They reasoned that the effect of these proteins was related to cell cycle arrest rather than cell death, based on the lack of cleaved caspase 3 in treated cells. This work focused on the role of this TP53-mediated response in homology-directed repair (HDR) and demonstrated that the addition of ectopic MDM2, a negative regulator of TP53, does increase the percentage of HDR-resolved cells in the assay. The authors interpreted this result as a specific effect on HDR, however an alternative explanation could be that MDM2 downregulation of TP53 simply allows the proliferation of the initial targeted cells, consistent with our observation of a Cas9-mediated arrest. The work of Ihry and colleagues (16) identified a TP53-dependent cytotoxicity phenotype in response to CRISPR/Cas9 treatment via differential expression analysis from RNA-seq data in human embryonic and induced pluripotent stem cells. An inducible version of DD-Cas9 was used in these experiments, and the assay identified p21 as the most differentially expressed gene. This is a surprising finding given the lack of increase of nuclear p21 in CRISPR/Cas9-treated RPE-1 cells in our experiments. While the identification of TP53 as the mediator in this cytotoxicity phenotype is interesting, this general phenotype has been previously observed (15).

A third recent study has proposed that CRISPR/Cas9 treatment only delays cell cycle progression (35). Interestingly, this delay was observed using a doxycycline-inducible, integrated version of DD-Cas9. As use of an inducibly-expressed, reversibly-stabilized Cas9 would be expected to limit the genomic occupancy time of Cas9, their results are consistent with our findings regarding DD-Cas9 and Shield-1 in addition to demonstrating a possible limit on the number of simultaneous DSBs that wild-type cells can recover from. This study examined a much earlier time point than our work (1 day as opposed to approximately 3 days post-transfection/induction) and relies on fluorescent reporters of cell cycle stage instead of incorporation of a nucleotide analog over an extended period of time, somewhat restricting the resolution of their analysis of cell cycle effect. This limitation is especially relevant in light of the observation that DNA damage in G1 can lead to arrest in the subsequent G2 phase (39). A conclusion of this paper (35) was that a small number of Cas9-induced DSBs do not lead to permanent cell cycle arrest, however, this must be viewed in light of the use of DD-Cas9 in their experiments, and that removal of DD-Cas9 from the locus would limit the inhibitory effect of Cas9, as we also observed. Thus, their results and ours support the preferential use of an inducible DD-Cas9 over wild-type for genome engineering applications.

The presence of [Cas9+ EdU+] wild-type RPE-1 cells in the experiments shown in Figure 1 suggests that these cells were able to progress through the cell cycle despite the Cas9 treatment. We noticed in analyzing our flow cytometry data that this population appears to be two subpopulations: (1) cells that have progressed through one cell cycle since onset of EdU incubation, (2) cells that have progressed through two cell cycles. Considering our experimental design, the first subpopulation most likely progressed through no more than two cell cycles before arresting, the first occurring sometime after transfection but before EdU incubation, and the second after EdU incubation. The majority of the second subpopulation most likely progressed through two cell cycles, beginning after the start of EdU incubation, before arresting after the second complete cell cycle. The remaining minority most likely reflects edited cells. The possibility that a substantial fraction of [Cas9+ EdU+] cells arrested after progressing through at least one cell cycle is supported by the recent finding that inducing DNA damage in G2 leads to cells arresting in the following G1 phase, whereas damage induced in G1 phase leads to arrest in the following G2 (39). Interestingly, that study also describes a diminished ability of U2-OS cells to permanently arrest in response to damage, which is consistent with our findings. This report also noted that damage in G1 results in a gradual switch from temporary to permanent arrest as the damage level increases, raising the possibility that persistent DNA damage, such as that induced by CRISPR/Cas9, might gradually lead to permanent arrest, which also could explain the [Cas9+ EdU+] fraction of cells.

In considering that Cas9 may occlude the DSBs it itself generates, one should also reflect on what occurs when using DD-Cas9. Once the stabilizing small molecule Shield-1 is withdrawn, DD-Cas9 becomes destabilized and susceptible to degradation. Thus, the DSB should then become available to repair. This availability appears to be reflected in the fact that the proportion of Cas9-cells increased in the population of DD-Cas9 transfected *TP53*^−/−^ RPE-1 cells (Supplementary Figure S3B), but not in transfected wild-type RPE-1 cells. However, availability of the DSB to repair is evident by the presence of nuclear 53BP1 foci in transfected wild-type RPE-1 cells (Figure 4C).

One major caveat of our work is that we chose to target a single non-transcribed locus. However, we would expect from recent work (33) that CRISPR/Cas9-associated arrest also occurs when targeting transcribed loci, with the degree of arrest related to the level of transcription through the locus. Similar TP53-dependent effects were observed in iPSCs when targeting transcribed loci (16), providing support for this being a general phenomenon in mammalian somatic cells. It is possible that embryos may be an exception due to their reliance for DNA damage repair on DNA Pol θ, which is relatively error-prone and might more frequently result in non-targetable products after an initial break (40). Going forward, determining how Cas9 is removed from DNA in the context of gene editing and how cells perceive Cas9-mediated double-strand breaks will aid in increasing the efficiency of genome engineering for experimental and therapeutic purposes.

## ACKNOWLEDGEMENTS

We thank the members of the Stearns lab for helpful and insightful commentary. Additionally, we thank the staff of the Stanford Shared FACS Facility, particularly Cathy Crumpton and Ometa Herman for sharing their expertise.

## FUNDING

This work was supported by funding from the National Institute of General Medicine (F32GM122214 to J.M.G.; R35GM130286 to T.S.).

## Supplementary Information

**Supplementary Figure 1.**
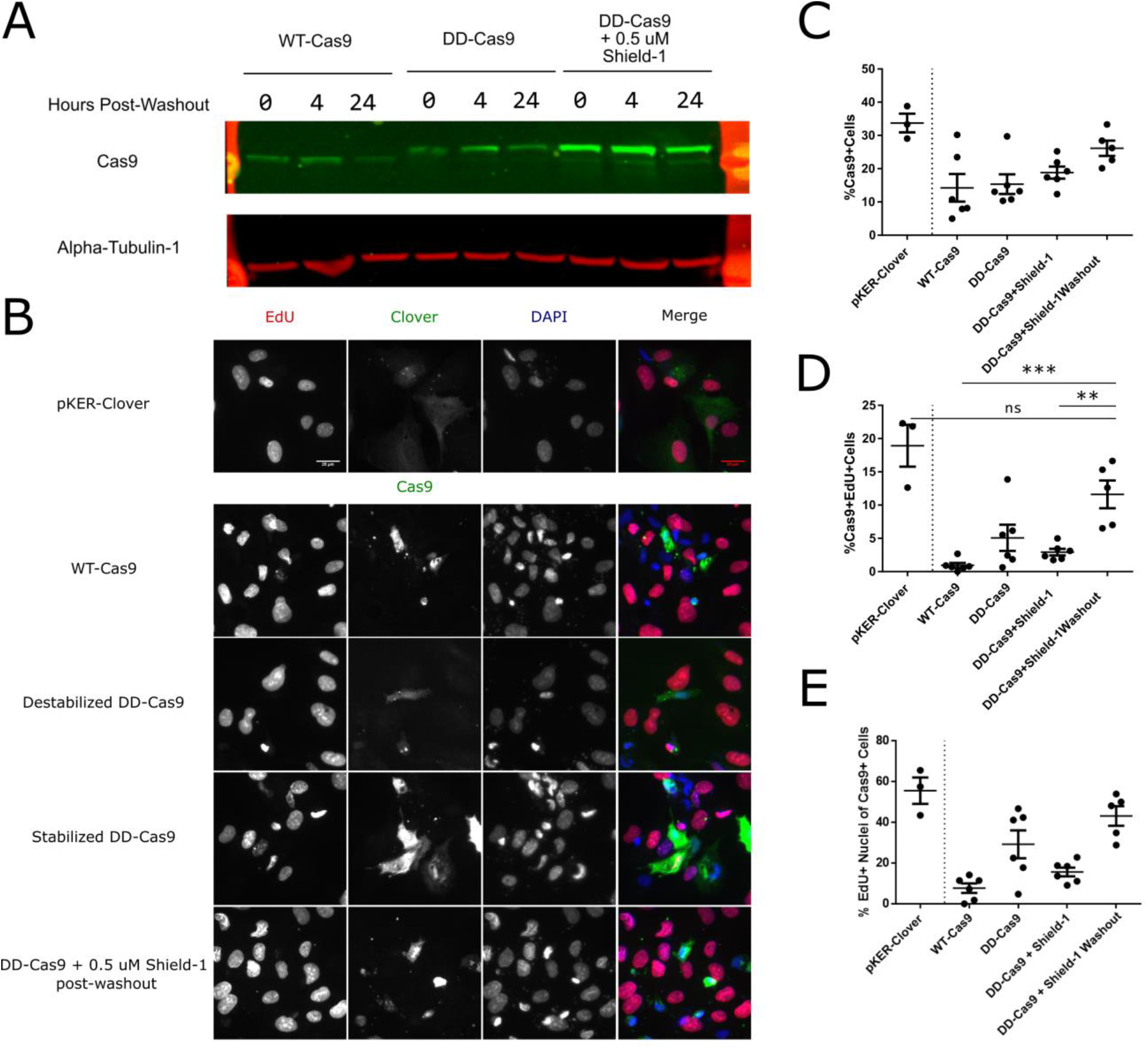
Wild-type SpCas9 decreases EdU incorporation in wild-type RPE-1 cells. (A) Western blot for wild-type Cas9 and DD-Cas9 in the absence or presence of 0.5 μM Shield-1 at the indicated times post-washout following a 24-hour incubation with 0.5 μM Shield-1. Green indicates Cas9 and red indicates alpha tubulin (used here as a loading control). (B) Immunofluorescence of RPE-1 cells transfected with the indicated vectors and incubated for roughly 45 hours in the presence of 10 μM EdU. Destabilized DD-Cas9 = no Shield-1. Stabilized DD-Cas9 = continuous Shield-1. Scale bar = 25 μm. (C) Dot plot of data from experiment described in (B) shown as percentage of transfected RPE-1 cells for indicated treatments. *n* = three technical replicates for pKER-Clover, two biological replicates each consisting of at least two technical replicates for all other transfections. Each dot is one technical replicate. (D) Dot plot of the Cas9+ EdU+ cells as a percentage of all cells counted from (B). (E) Dot plot of the EdU+ cells depicted in (D) as a percentage of transfected cells (Cas9+). Data were analyzed with a one-way ANOVA followed by a post hoc Tukey’s multiple comparisons test. Error bars indicate SEM. ** = *P* < 0.005. *** = *P* = 0.0001. ns = not significant.

**Supplementary Figure 2.**
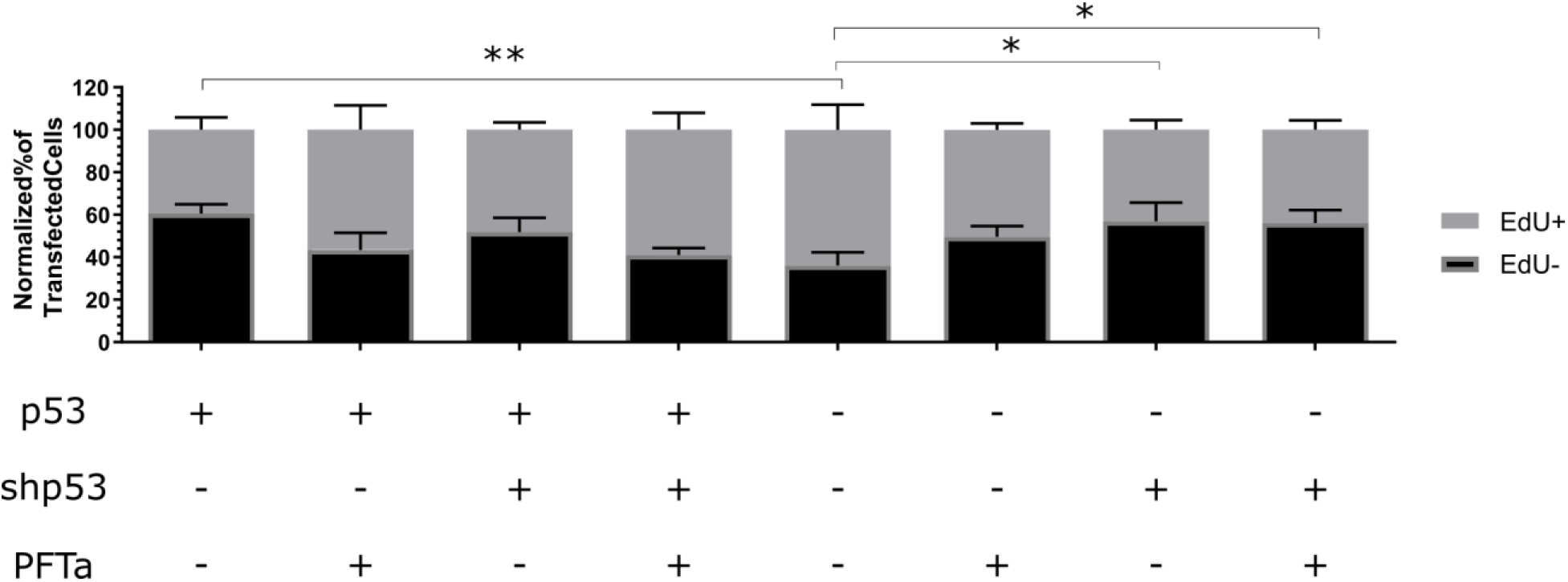
*TP53* RNAi negatively impacts cell cycle progression in *TP53*^−/−^ RPE-1 cells. Bar graph of Cas9+ normalized percentages of EdU+ and EdU-cells for indicated treatments. *n* = at least two independent experiments each consisting of three technical replicates for each transfection. Data is shown as the mean ± SEM. Data were analyzed with a two-way ANOVA followed by a post hoc Dunnet’s multiple comparisons test using *TP53*^−/−^ RPE-1 + WT-Cas9 as the control sample. * = *P* < 0.05. ** = *P* < 0.01.

**Figure S3.**
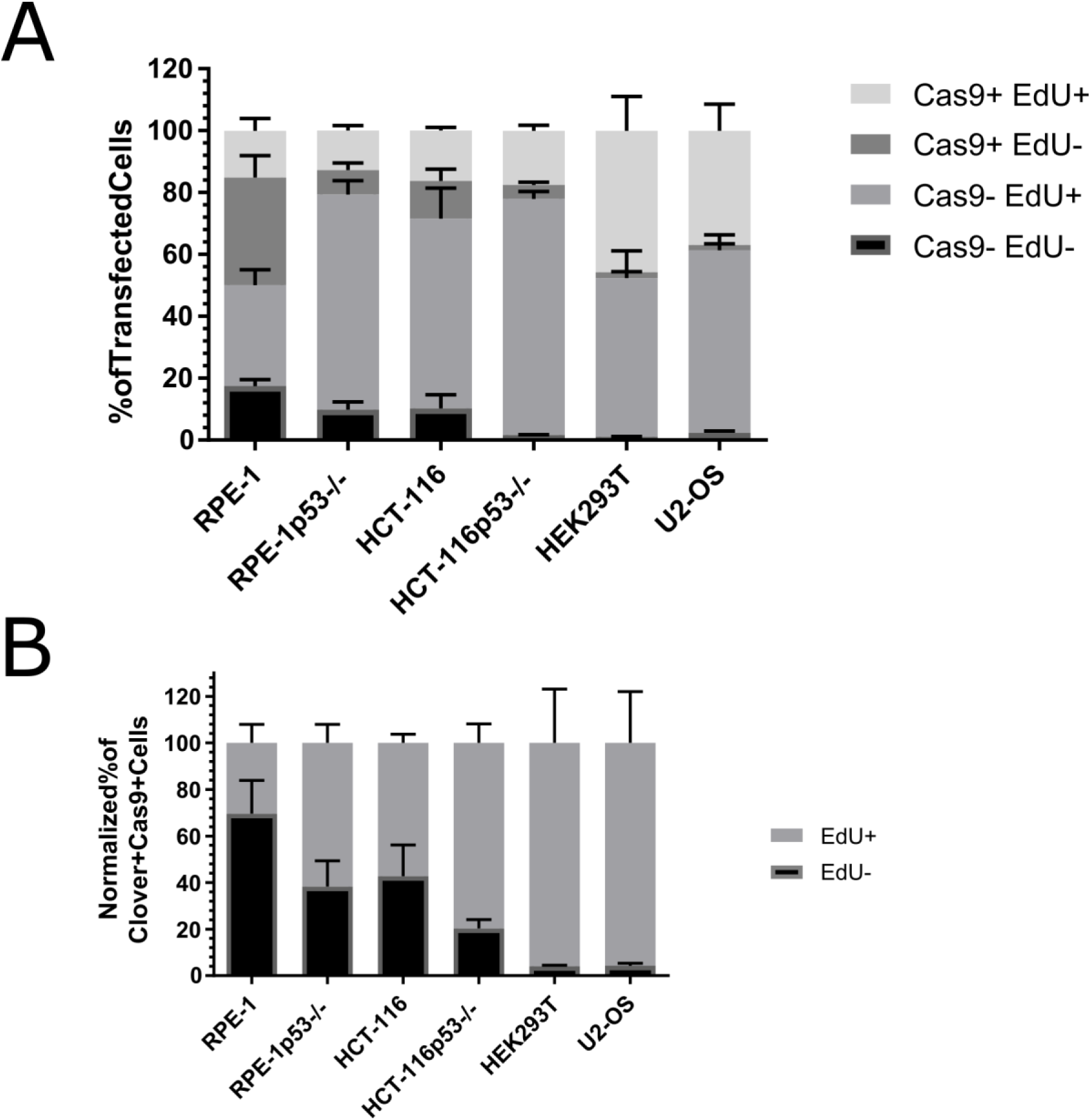
Distribution of cell cycle progression in cell lines treated with temporarily stabilized DD-Cas9. (A) Bar graph of the normalized percentages of four Cas9 EdU subpopulations in transfected cells for the six indicated cell lines. *n* = at least two independent experiments consisting of three technical replicates for each transfection. Data are shown as the mean ± SEM. (B) Bar graph of the Cas9+ normalized percentages of EdU+ and EdU-from panel A. Data is shown as the mean ± SEM.

